# Karyological diversity among six medicinally important species of Acanthaceae in Bangladesh

**DOI:** 10.1101/2023.07.02.547390

**Authors:** Nowshin Anjum, Chandan Kumar Dash, Syeda Sharmeen Sultana

## Abstract

Six medicinally important species of Acanthaceae family namely, *Asystasia gangetica* (L.) T. Anderson, *Barleria cristata* L., *Braleria prionitis* L., *Ruellia simplex* C. Wright, *Thunbergia erecta* (Benth.) T. Anderson and *Thunbergia mysorensis* (Wright) T. Anderson were cytogenetically investigated to evaluate the karyotype diversity and chromosomal relationship. The somatic chromosome numbers and the centromeric formulas of these species were 2n = 2x = 24 (22m + 2sm) for *A. gangetica*, 2n = 4x = 40 (38m + 2sm) for *B. cristata*, 2n = 4x = 40 (36m + 4sm) for *B. prionitis*, 2n = 2x = 36 (34m + 2sm) for *R. simplex*, 2n = 4x+6 = 62 (56m + 6sm) for *T. erecta* and 2n = 2x = 34 (34m) for *T. mysorensis*. Considering the karyotype assessment parameters, *T. erecta, R. simplex, B. cristata* and *B. prionitis* were shown to be cytogenetially more advanced than *A. gangetica* and *T. mysorensis*. Three categories of karyotypes were observed such as 1B type in *B. cristata, B. prionitis, R. simplex* and *T. mysorensis*, whereas the most primitive one 1A in *A. gangetica* and the most advanced karyotype 2B in *T. erecta*. Three new reports of somatic chromosome count yielded in case of *A. gangetica, R. simplex* and *T. mysorensis* which might have been resulted from aneuploidy or allopolyploidy. The cytological data presented for six species of the Acanthaceae family demonstrated significant variation and distinctiveness from one another, assisting in the evaluation of the karyological diversity that aided in the understanding of evolutionary changes.

## Introduction

The family Acanthaceae consists of approximately 240 genera and 3250 species of herbs and shrubs distributed throughout the world (Wasshausen and Wood 2004). Indo-Malaya, Africa, Brazil and Central America are the four main centers of distribution of this family (Cramer 1998). So far, 39 genera and 107 species under the family Acanthaceae have been reported from Bangladesh (Ahmed et al. 2008). Among these species *Asystasia gangetica* (L.) T. Anderson, *Barleria cristata* L., *Braleria prionitis* L., *Ruellia simplex* C. Wright, *Thunbergia erecta* (Benth.) T. Anderson and *Thunbergia mysorensis* (Wright) T. Anderson are most widely distributed in Bangladesh with promising medicinal and ecological values (Ahmed et al. 2008). These plants showed the presence of important secondary metabolites such as alkaloids, steroidal aglycones, saponins, flavonoids, reducing sugars, triterpenoids and have been claimed to have anti-inflammatory, anti-diabetic, anti-asthmatic, analgesic and broncho-spasmolytic properties (Gambhire et al. 2009; Singh and Kimothi 2018; Begum et al. 2019). These plants are also used for the treatment of toothache, anemia, snake bite, diabetes, lungs disorders, gastrointestinal disorders, blood diseases, boils, glandular swellings, ulcers, greying of hairs, fever, cough, jaundice, liver diseases, arthritis, etc. (Akah et al. 2003; Kale et al. 2003; Chen et al. 2006; Daniel 2006; Khare 2007; Gambhire et al. 2009; Singh and Kimothi 2018; Begum et al. 2019).

According to previous literature, in the case of *A. gangetica*, 2n = 26 was observed in most of the studies where the basic chromosome number was x = 13 (Gadella 1977; Ugborogho and Adetula 1988; Pandit et al. 2006). In addition, 2n = 25 (Narayanan 195l; Kumar and Subramaniam 1987), 2n = 28 (Subramanian and Govindarajan 1980, Govindarajan and Subramanian 1983) and 2n = 44, 48, 50 and 52 (Narayanan 1951; Grant 1955; Ellis 1962; Kaur 1965; Sarkar et al. 1978; Kumar and Subramaniam 1987) were reported by different scientists (Table 1). A wide range of somatic chromosome number for *B. cristata* had been reported such as 2n = 32, 34, 35, 38, 40 and 42 (Narayanan 1951; Datta and Maiti 1970; Saggo and Bir 1986; Kumar and Subramaniam 1987; Devi and Mathew 1991). *Braleria prionitis* has comparatively less chromosome number variation than *B. cristata* as 2n = 32 and 40 were reported in the previous literature (Datta and Maiti 1970; Krishnaswami and Menon 1974; Moore 1977; Saggo and Bir 1986; Devi and Mathew 1991; Shendage 2022). Only 2n = 34 had been confirmed as somatic chromosome number of *R. simplex* (Grant 1955; Long 1964; De 1966; Bedi et al. 1981; Saggoo 1983; Daniel and Chuang 1998). *Thunbergia erecta* has wide range of variation in somatic chromosome number. 2n = 16, 52, 56, 60, 62 and 64 were reported previously (Narayanan 1951; Grant 1955; Mangenot and Mangenot 1957; Mangenot and Mangenot 1962; Kaur 1969; Sareen and Sanjogta 1976; Subramanian and Govindarajan 1980). Goldblatt and Jhonson (1981) reported 2n = 28 chromosomes in *T. mysorensis* (Table 1).

**Table 1.**
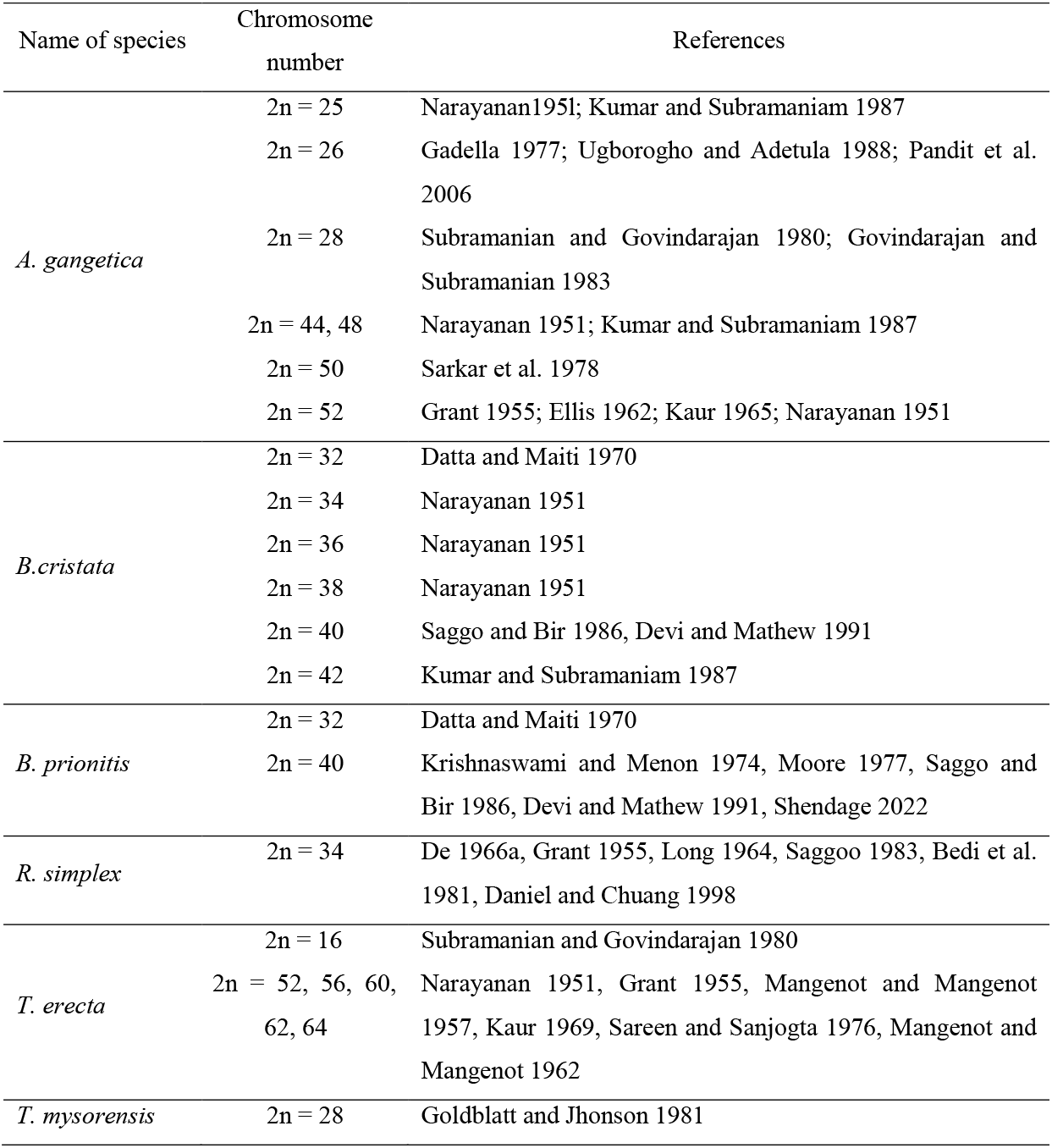
Previous chromosome number records of six species of Acanthaceae family.

| Name of species | Chromosome number | References |
| --- | --- | --- |
| *A. gangetica* | 2n = 25 | Narayanan 1951; Kumar and Subramaniam 1987 |
|  | 2n = 26 | Gadella 1977; Ugborogho and Adetula 1988; Pandit et al. 2006 |
|  | 2n = 28 | Subramanian and Govindarajan 1980; Govindarajan and Subramanian 1983 |
|  | 2n = 44, 48 | Narayanan 1951; Kumar and Subramaniam 1987 |
|  | 2n = 50 | Sarkar et al. 1978 |
|  | 2n = 52 | Grant 1955; Ellis 1962; Kaur 1965; Narayanan 1951 |
| *B. cristata* | 2n = 32 | Datta and Maiti 1970 |
|  | 2n = 34 | Narayanan 1951 |
|  | 2n = 36 | Narayanan 1951 |
|  | 2n = 38 | Narayanan 1951 |
|  | 2n = 40 | Saggo and Bir 1986, Devi and Mathew 1991 |
|  | 2n = 42 | Kumar and Subramaniam 1987 |
| *B. prionitis* | 2n = 32 | Datta and Maiti 1970 |
|  | 2n = 40 | Krishnaswami and Menon 1974, Moore 1977, Saggo and Bir 1986, Devi and Mathew 1991, Shendage 2022 |
| *R. simplex* | 2n = 34 | De 1966a, Grant 1955, Long 1964, Saggoo 1983, Bedi et al. 1981, Daniel and Chuang 1998 |
| *T. erecta* | 2n = 16 | Subramanian and Govindarajan 1980 |
|  | 2n = 52, 56, 60, 62, 64 | Narayanan 1951, Grant 1955, Mangenot and Mangenot 1957, Kaur 1969, Sareen and Sanjogta 1976, Mangenot and Mangenot 1962 |
| *T. mysorensis* | 2n = 28 | Goldblatt and Jhonson 1981 |

Consequently, karyotype might be viewed as a wonderful source of data for evolutionary research among various species (Stace 2000; Dobigny et al. 2004; Guerra 2012). In Bangladesh, the chromosome analysis of these six important species under the family Acanthaceae has not yet been initiated. The diversity regarding somatic chromosome counts and the multi-basic nature make those species attractive for cytogenetical research. Karyological data can be utilized to assess the taxonomic relationship among taxa for better understanding the evolutionary relationship and diversity (Guerra 2008). Therefore, in this present study, a detailed karyological analysis of six species of the Acanthaceae family was carried out to evaluate the chromosomal relationships among them with morphological implications regarding the evolutionary perspective.

## Materials and Methods

The plant materials of this investigation, *Asystasia gangetica* (L.) T. Anderson, *Barleria cristata* L., *Braleria prionitis* L., *Ruellia simplex* C. Wright, *Thunbergia erecta* (Benth.) T. Anderson and *Thunbergia mysorensis* (Wright) T. Anderson were collected from different nurseries of Dhaka city and Dhaka University campus. The experimental plots were maintained in the Botanical Garden, Department of Botany, University of Dhaka.

The root tips were collected and soaked on a filter paper to remove surface water and kept in 8-hydroxyquinoline (0.002 M) for 4 h at 10–12 °C for pre-treatment. The pre-treated roots were removed from 8-hydroxyquinoline (0.002 M) and preserved in Carnoy’s solution (Absolute alcohol: Glacial acetic acid = 3: 1) at 4 °C. The pretreated RTs were hydrolyzed for 8 min at 65 °C in a mixture of 1N HCl and 45% acetic acid (2:1). Then the hydrolyzed RTs were soaked on a filter paper and taken on a clean oil free slide. The meristematic region, which is a small portion of the root tip, was cut with a fine blade. A drop of 1% aceto-orcein was added to the material and kept in an acetic acid chamber for 2 h 30 min. Then, a clean cover glass was placed on the material. At first, the materials were tapped gently by a toothpick and then squashed by placing thumbs. The slides were observed under Euromax Axion microscope. To get an accurate measurement of lengths, chromosomes from five metaphase plates were measured for each species. KaryoType software was used to measure the length of the chromosomes (Altinordu et al. 2016). The average arm length was used to prepare the karyotype. The chromosomes were arranged gradually from longer to shorter in length. The short arm was placed on the upper side of the axis and the long arm on the lower side. The haploid idiograms were prepared with the help of KaryoType software to represent the karyotypes more clearly (Altinordu et al. 2016). The box plot was prepared by R-programming. The method by Levan et al. (1964) was used to determine the centromeric type of chromosomes. Based on Stebbins’ (1971) classification, karyotypes were categorized. KaryoType software (Altinordu et al. 2016) was used to calculate a variety of karyotypic symmetric and asymmetric indices including total form percent (TF %) by Huziwara (1962), karyotype asymmetry index (AsK%) by Arano (1963), chromosome size resemblance index (Rec%) and karyotype symmetry index (Syi%) by Greilhuber and Speta (1976), intrachromosomal asymmetry index (A_1_) and interchromosomal asymmetry index (A_2_) by Zarco (1986), dispersion index (DI) of Lavania and Srivastava (1992), degree of karyotype asymmetry (A) by Watanabe et al. (1999), asymmetry index (AI), coefficient of variation of the centromeric index (CV_CI_) and coefficient of variation of the chromosome length (CV_CL_) by Paszko (2006) and mean centromeric asymmetry (M_CA_) by Peruzzi and Eroğlu (2013). For the cluster analysis, we used the UPGMA algorithm (unweighted pair-group method with arithmetic means) implemented in the online software DendroUPGMA, http://genomes.urv.cat/UPGMA/.

## Results

The morphological descriptions such as habit, plant’s height, leaf type and arrangement, leaf size, leaf shape, number of stamen, petal and sepal, inflorescence type, flowering and fruiting time, flower colour, fruit types and number of seeds per fruit of six Aanthaceae species are provided in table 2. The results regarding the extensive karyological analysis such as somatic chromosome number, total chromosomal length, average chromosomal length, range of chromosomal length and different symmetric and asymmetric karyotypic indices are presented in table 3.

**Table 2.** Morphological description of six species from Acanthaceae family.

| Plant<br>Features | *A. gangetica* | *B. cristata* | *B. prionitis* | *R. simplex* | *T. erecta* | *T. mysorensis* |
| --- | --- | --- | --- | --- | --- | --- |
| Habit | Climbing sub-shrub | Shrub | Shrub | Erect herb | Shrub | Climber |
| Plant's height | Up to 0.5 m | Up to 2 m | Up to 1.8 m | 30-50 cm | Up to 2 m | Up to 6 m |
| Leaf type and arrangement | Simple, opposite | Simple, opposite | Simple, opposite | Simple, opposite | Simple, opposite | Simple, opposite |
| Leaf Size | ~2 cm×1–2 cm | 2-10×1–4 cm | 4–10.5×1.8–5.5 cm | 6–12 inches | 4–8×2.0–3.5 cm | 10–15 cm |
| Leaf Shape | Elliptic, ovate | Elliptic-oblong to lanceolate | Elliptic to ovate | Leaves lanceolate, narrowed at both ends, serrate | Ovet- elliptic to obovate, serrate, acute | Elliptic or oblong lanceolate |
| Stamen | Four, included | Four, didynamous | Four, didynamous | Four, didynamous | Four, didynamous | Four, didynamous |
| Petal and sepal | Calyx 5-lobed; Corolla funnel-shaped, 5-lobed | Calyx deeply 4-lobed; Corolla funnel-shaped, 5-lobed | Calyx deeply 4-lobed; Corolla funnel-shaped, 5-lobed | Calyx deeply 5-lobed; Corolla funnel-shaped, 5-lobed | Calyx 15 toothed; Corolla funnel-shaped, 5-lobed | Calyx 15 toothed; Corolla funnel-shaped, 5-lobed |
| Inflorescence | Axillary racemes | short and dense cymes | Cyme | Axillary cyme | Cyme | Racemes |
| Flowering and fruiting time | September to February | November to February | November to February | July to October | October to January | January to March |
| Flower colour | White | White | Yellow | Violet-blue | Blue | Brownish red with yellow |
| Fruit | Capsule | Capsule | Capsule | Capsule | Capsule | Capsule |
| Seeds per fruit | up to 4 seeded | 4 seeded | 2 seeded | 4-28 seeded | 2–4 seeded | 2–4 seeded |

**Table 3.** Comparative karyological analysis of six species of Acanthaceae family.

| Species<br>Features | *A. gangetica* | *B. cristata* | *B. prionitis* | *R. simplex* | *T. erecta* | *T. mysorensis* |
| --- | --- | --- | --- | --- | --- | --- |
| 2n | 24 | 40 | 40 | 36 | 62 | 34 |
| CF | 22m + 2sm | 38m + 2sm | 36m + 4sm | 34m + 2sm | 56m + 6sm | 34 m |
| TCL (μm) | 33.71 | 92.76 | 112.81 | 69.73 | 107.72 | 58.93 |
| ACL (μm) | 1.40 | 2.32 | 2.82 | 1.94 | 1.74 | 1.73 |
| RCL (μm) | 1.17 – 1.77 | 1.29 – 3.72 | 1.88 – 4.25 | 1.16 – 3.28 | 0.65 – 2.55 | 1.08 – 2.88 |
| CV<sub>CI</sub> | 7.10 | 6.16 | 6.48 | 7.73 | 8.65 | 3.78 |
| CV<sub>CL</sub> | 12.64 | 26.21 | 21.38 | 23.92 | 24.84 | 19.05 |
| M<sub>CA</sub> | 8.16 | 6.46 | 9.91 | 6.69 | 6.31 | 5.99 |
| AsK % | 54.26 | 53.21 | 54.79 | 53.33 | 53.59 | 53.14 |
| TF % | 45.74 | 46.79 | 45.21 | 46.34 | 46.41 | 46.86 |
| Syi % | 84.30 | 87.93 | 82.50 | 86.38 | 86.60 | 88.17 |
| Rec % | 79.31 | 62.35 | 66.29 | 58.90 | 68.15 | 61.17 |
| A<sub>1</sub> | 0.15 | 0.12 | 0.18 | 0.12 | 0.11 | 0.11 |
| A<sub>2</sub> | 0.13 | 0.26 | 0.21 | 0.24 | 0.25 | 0.19 |
| A | 0.08 | 0.06 | 0.10 | 0.07 | 0.06 | 0.06 |
| AI | 0.90 | 1.61 | 1.39 | 1.85 | 2.15 | 0.72 |
| DI | 5.89 | 12.28 | 9.85 | 11.13 | 11.35 | 0.89 |
| Category | 1A | 1B | 1B | 1B | 2 B | 1B |

[2n = Somatic chromosome number, CF = Centromeric formula, TCL = Total chromosome length (μm), ACL = Average chromosome length (μm), RCL = Range of chromosomal length, CV<sub>CI</sub> = Coefficient of variation of centromeric index, CV<sub>CL</sub> = Coefficient of variation of chromosome length, M<sub>CA</sub> = Mean centromeric asymmetry, AsK% = Karyotype asymmetry index (%), TF% = Total form value (%), Syi% = Karyotype symmetry index (%), Rec% = The index of chromosomal size resemblance, A<sub>1</sub> = Intrachromosomal asymmetry index, A<sub>2</sub> = Interchromosomal asymmetry index, A = Degree of asymmetry of karyotypes, AI = The asymmetry index, DI = The dispersion index, m = metacentric chromosome, sm = sub-metacentric chromosome].

### Chromosome number and karyotype formula

*Asystasia gangetica* was found to possess 2n = 24 chromosomes (Figure 1a). The karyotypic formula was 22m + 2sm (Figure 2a, Table 3). The chromosomal length ranged from 1.17 – 1.77 μm and the total chromosome length was 33.71 μm. The average chromosome length for this species was 1.40 μm (Table 3). The somatic chromosome was found to be 2n = 40 in both *B. cristata* and *B. prionitis* (Figures 1b, 1c). However, they differed in terms of karyotypic formula i.e., 38m + 2sm for *B. cristata* and 36m + 4sm for *B. prionitis* (Figures 2b, 2c). The total and average chromosomal length was 92.76 μm and 2.32 μm in *B. cristata* and 112.81 μm and 2.82 μm in *B. prionitis*, sequentially. The range of longest and shortest chromosomes was 1.29 – 3.72 μm in *B. cristata* and 1.88 – 4.25 μm in *B. prionitis* (Table 3). 2n = 36 chromosomes were observed in *R. simplex* in which 34 chromosomes were metacentric and 2 were submetacentric (Figures 1d, 2d). The length of chromosomes ranged from 1.16 – 3.28 μm. The total and average chromosomal length was 69.73 μm and 1.94 μm, respectively (Table 3). The somatic chromosome count was 2n = 62 for *T. erecta* and 2n = 34 for *T. mysorensis* (Figure 1e, 1f). The karyotype formula of *T. erecta* was 56m + 6sm whereas all the chromosomes were metacentric in *T. mysorensis* (Figure 2e, 2f, Table 3, for details, see supplementary tables).

**Figure 1.**
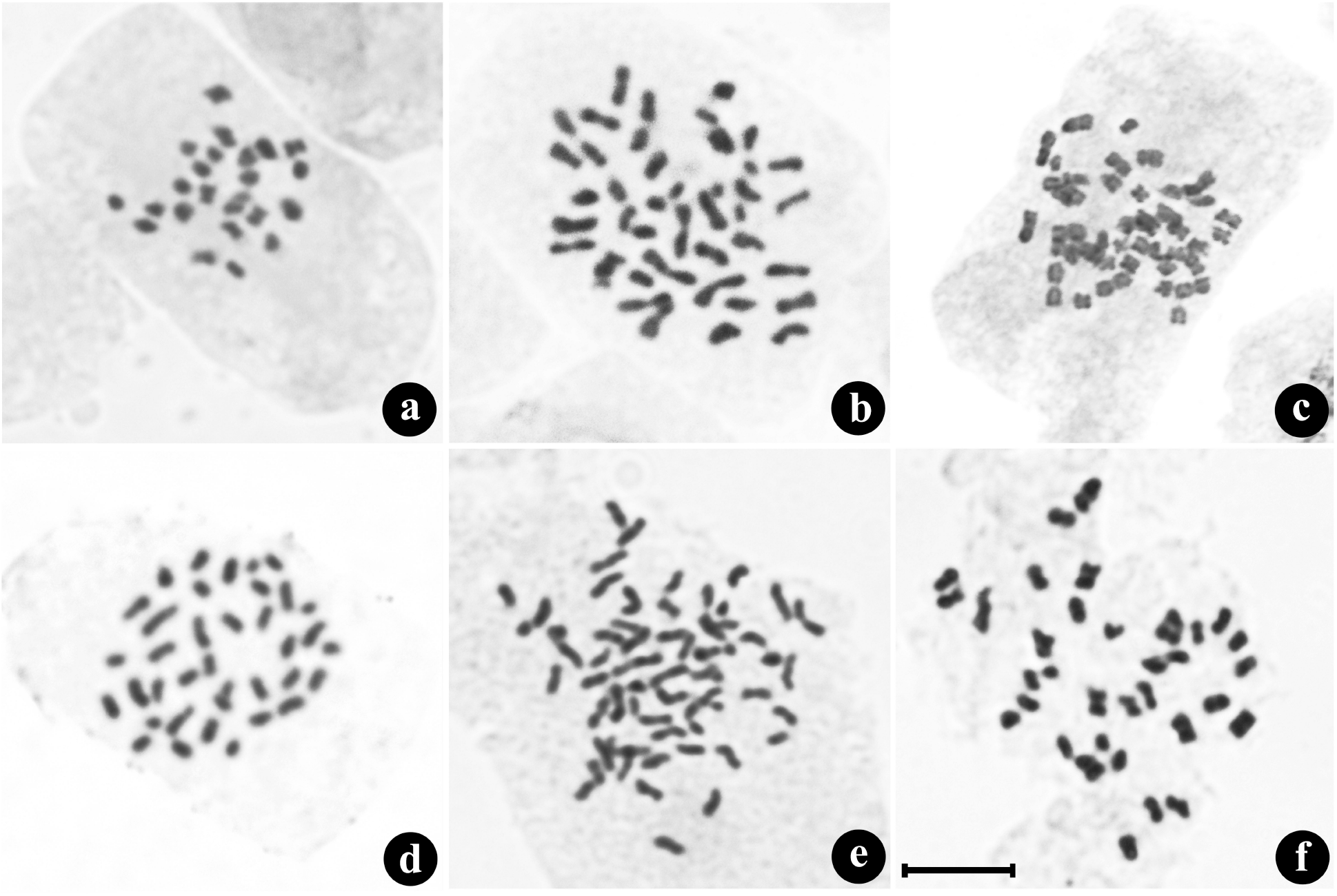
Orcein-stained metaphase chromosomes of six species of Acanthaceae family; (a) *Asystasia gangetica*; (b) *Barleria cristata*; (c) *Braleria prionitis*; (d) *Ruellia simplex*; (e) *Thunbergia erecta*; (f) *Thunbergia mysorensis* Scale bar = 10 μm.

**Figure 2.**
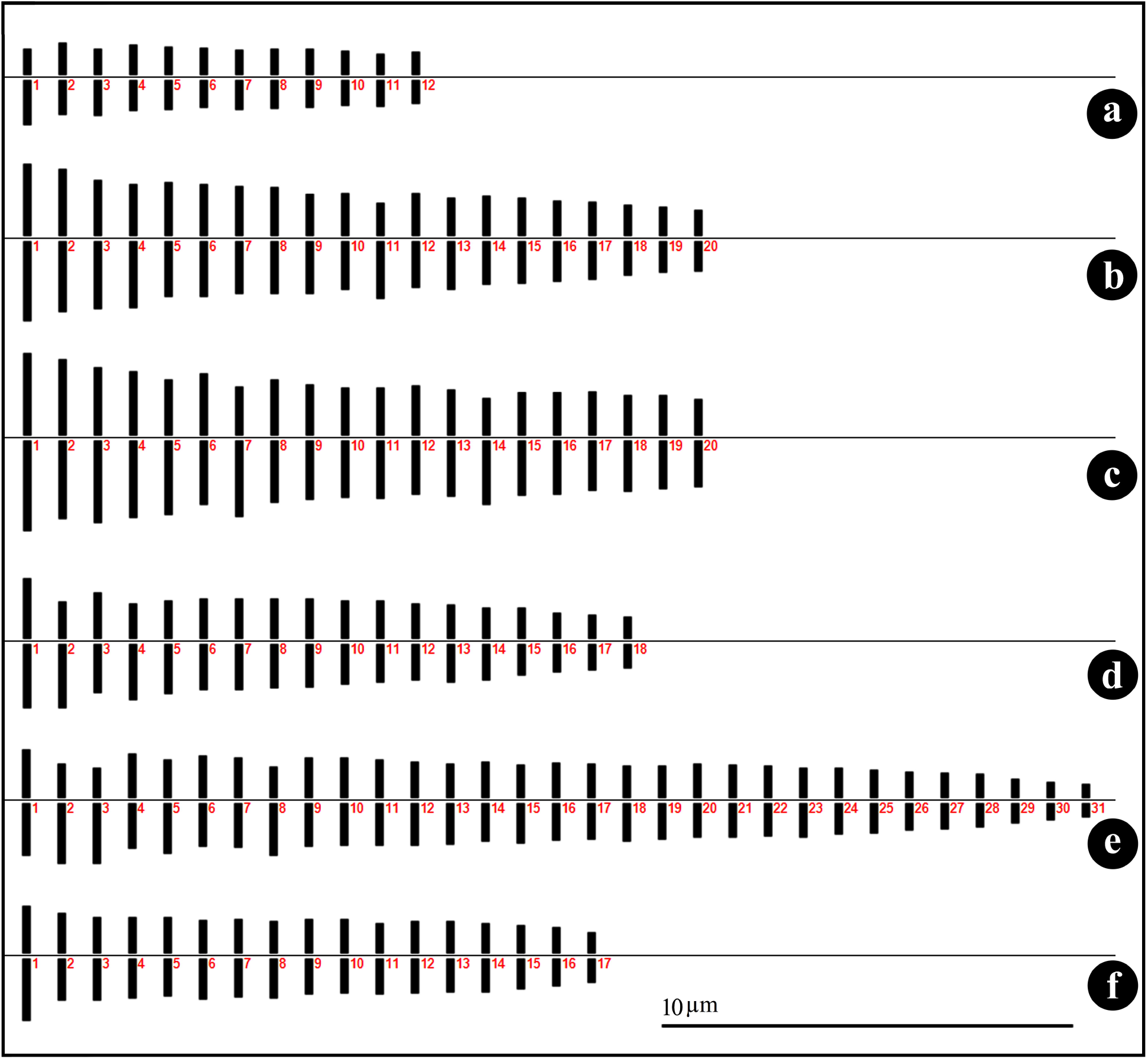
Idiograms of six species of Acanthaceae family; (a) *Asystasia gangetica*; (b) *Barleria cristata*; (c) *Braleria prionitis*; (d) *Ruellia simplex*; (e) *Thunbergia erecta*; (f) *Thunbergia mysorensis* Scale bar = 10 μm.

### Assessment of symmetric and asymmetric karyotypic indices

The CV_CI_ values for *A. gangetica, B. cristata, B. prionitis, R. simplex, T. erecta* and *T. mysorensis* were 7.10, 6.16, 6.48, 7.73, 8.65 and 3.78, whereas the CV_CL_ values were 12.64, 26.21, 21.38, 23.92, 24.84 and 19.05, respectively. The asymmetry index (AI) was higher in *T. erecta* (2.15) and relatively lower in *R. simplex* (1.85), *B. cristata* (1.61), *B. prionitis* (1.39), *A. gangetica* (0.90) and *T. mysorensis* (0.72) (Table 3). All the studied species were in the same category (1B) except *A. gangetica* (1A) and *T. erecta* (2B) according to Stebbins (1971). The karyotype asymmetry (AsK%) index was highest in *B. prionitis* (54.79%) and lowest in *T. mysorensis* (53.14%). In case of karyotype symmetry (Syi%) index, the value was found lowest in *B. prionitis* (82.50%) and highest in *T. mysorensis* (88.17%). The mean centromeric asymmetry (M_CA_) in *A. gangetica, B. cristata, B. prionitis, R. simplex, T. erecta* and *T. mysorensis* were observed as 8.16, 6.46, 9.91, 6.69, 6.31 and 5.99 and total form value (TF%) was 45.74, 46.79, 45.21, 46.34, 46.41 and 46.86, sequentially. The index of chromosomal size resemblance (Rec%) was highest in *A. gangetica* (79.31), whereas *B. cristata, B. prionitis, R. simplex, T. erecta* and *T. mysorensis* showed 62.35, 66.29, 58.90, 68.15 and 61.17, accordingly. However, A_1_, A_2_, A and DI values for *A. gangetica* (0.15, 0.13, 0.08, 5.89), *B. cristata* (0.12, 0.26, 0.06, 12.28), *B. prionitis* (0.18, 0.21, 0.10, 9.85), *R. simplex* (0.12, 0.24, 0.07, 11.13), *T. erecta* (0.11, 0.25, 0.06, 11.35) and *T. mysorensis* (0.11, 0.19, 0.06, 8.89) were recorded, respectively (Table 3).

## Discussion

### New chromosome count and karyotype for Asystasia gangetica

*Asystasia gangetica* was found to possess 2n = 24 chromosomes which is a new chromosome count (Figures 1a, 2a, Table 3). Previously, all the cytological reports regarding this species were confined mostly to chromosome counting and no karyotype has been reported so far except the report of Govindarajan and Subramanian (1983). In this observation, 22 metacentric along with 2 submetacentric chromosomes were found with the range of 1.17 – 1.77 μm (Table 3). Different scientists mentioned 2n = 25 for this species (Narayanan 195l; Mangenot and Mangenot 1957, 1962; Kumar and Subramaniam 1987). Previously published chromosomal reports showed that mostly 2n = 26 was observed with basic chromosome number x = 13 (Gadella 1977; Ugborogho and Adetula 1988; Daniel 2000; Pandit et al. 2006) which was not observed in the present study (Table 1). Pandit et al. (2006) reported 13 ring bivalents in most of the cases at meiotic metaphase I. 2n = 28 for this species might be found due to hyper-aneuploidy or the presence of B-chromosome as observed by Govindarajan and Subramanian (1983) and Subramanian and Govindarajan (1980). 2n = 50 (Sarkar et al. 1978) and 2n = 44, 48 (Narayanan 195l; Grant 1955; Kumar and Subramaniam 1987) were also observed. The presence of 2n = 4x = 52 strongly supported the basic chromosome number x = 13 (Narayanan 1951; Grant 1955; Ellis 1962; Kaur 1965). Saggoo and Bir (1986), on the other hand, suggested four alternative basic chromosome numbers for *Asystasia*, x = 11, 12, 13 and 16. According to earlier reports, we can consider x = 13 to be the most predominantly basic chromosome number for *Asystasia* and it is possible that it was caused by aneuploidy or allopolyploidy. Pandit et al. (2006) hypothesized that the hybridization of two-parent genomes, one with 2n = 12 and another with 2n = 14, gave rise to the hybrid genome (2n = 13) and further genome duplication may have resulted in an allopolyploid with 2n = 26. However, 2n = 24 for *A. gangetica* in the present study indicated the probable occurrence of new basic x = 12 as they showed normal vegetative growth with viable seed rather considered as aneuploidy (2n – 2 = 24). Whatever the reason, 2n = 24 is the new report for *A. gangetica* from Bangladesh.

### Ploidy status and karyomorphology of two Barleria species

In the present study, 2n = 40 chromosomes were found in *B. cristata* and *B. prionitis* (Figures 1b, 1c, 2b, 2c) which correlates with most of the previous findings (Krishnaswami and Menon 1974; Moore 1977; Saggo and Bir 1986; Devi and Mathew 1991; Shendage 2022). In this context, *B. cristata* and *B. prionitis* could be regarded as tetraploid (2n = 4x = 40). *Barleria* possesses a less diversified chromosome count according to previous reports (Table 1). Somatic counts that deviate from 2n = 40 such as 2n = 30, 32, 34, 36, 38 and 42 have been attributed to aneuploid changes and hybridization in several horticultural cultivars of *Barleria cristata* (Ranganath and Krishnappa 1990). Shendage (2022) reported 16 large and 24 relatively small metacentric chromosomes in *B. prionitis*, and categorized as 2B according to Stebbins’s (1971) classification. In this present investigation, both *Barleria* showed 1B karyotype which was not supported by the previous findings. According to Shendage (2022), the length of haploid chromosome complements of *B. prionitis* was 72.35 μm. Here, the total length of diploid chromosome complements of *B. prionitis* was 112.81 μm with average length of 2.82 μm (Table 3). The variation of chromosomal length may be due to the effect of pretreating agents and time.

### 2n = 36 is a new report for Ruellia simplex

In this investigation, the studied *R. simplex* possess 2n = 36 which is different from most of the previous literature indicating the probable occurrence of cytotype (Figures 1d, 2d, Table 3). Several scientists found diploid chromosome number 2n = 34 with the most frequent basic chromosome number x = 17 for this species (Grant 1955; Long 1964; De 1966; Bedi et al. 1981; Saggoo 1983; Daniel and Chuang 1998). The diversity in chromosome number can be explained either by non-disjunction or possible hybridization within different cytotypes among these species belonging to the same genus. Grant (1955) reported 2n = 34 chromosome numbers for 27 taxa under the genus *Ruellia*, as well as a few unusual counts of n = 16 (*R. cordata*) (Rao and Mwasumbi 1981). Additionally, different scientists reported 2n = 32 in *R. patula* and *R. tuberosa*, 2n = 36 in *R. amoena, R. ciliate* and *R. dipteracamthus* (Fedorov 1974; Govindarajan and Subramanian 1983) which revealed the occurrence of multiple basic chromosome numbers x = 16 and 18 along with the most common x = 17.

### Variable chromosome number in Thunbergia genus with new count for T. mysorensis

In this investigation, *Thunbergia* L. species showed variation in somatic chromosome counts such as 2n = 62 in *T. erecta* and 2n = 34 in *T. mysorensis* with different basic chromosome numbers (Figures 1e, 1f, 2e, 2f, Table 3). According to the previous chromosome number records, 2n = 16 (Govindarajan and Subramanian 1983) and 2n = 52, 56, 60, 62 (Narayanan 1951; Grant 1955; Mangenot and Mangenot 1957; Kaur 1969; Sareen and Sanjogta 1976; Mangenot and Mangenot 1962) were reported for *T. erecta*. Grant (1955) proposed x = 7 as the actual basic chromosome number for the Acanthaceae family. Later, Long (1970) hypothesized that the lowest haploid number for this family observed in *Thunbergia*, i.e., n = 7 could be Acanthaceae’s basic primitive number, from which a series of basic numbers along with the most common n = 14 could have been originated (Piovano and Bernardello 1991). According to Saggoo and Bir (1982), *Thunbergia* represents multibasic chromosome numbers such as x = 9, 10, 14, 16. The present findings (2n = 62) supported the previous report of Sareen and Sanjogta (1976), and the ploidy level could have been 4x + 6. Sareen and Sanjogta (1976) did not explain the probable cause of that hyper-tetraploidy. Cytogenetically *T. mysorensis* was very less studied and there was no report on the chromosome count except 2n = 28 (Goldblatt and Jhonson 1981) which was not supported by the present finding 2n = 34. Therefore, 2n = 34 is the new report of *T. mysorensis* (Figure 1f, Table 3). Therefore, the chromosome number variation for this genus might be originated from secondary modification of polyploids or hybridization among different cytotypes. Unstable and variable chromosome count in this genus suggested that aneuploidy and polyploidy played an important role in the evolution of a series of new basic chromosome numbers, accompanied by the diversification of species within the genus *Thunbergia*.

### Karyotype diversity among six species

Predominantly all the studied species possess metacentric chromosomes with few submetacentric chromosomes and the centromeric formulae are 22m + 2sm, 38m + 2sm, 36m + 4sm, 34 m + 2sm, 56m + 6sm and 34m for *A. gangetica, B. cristata, B. prionitis, R. simplex, T. erecta*, and *T. mysorensis*, respectively (Table 3). *T. erecta* has more heterogenous karyotype due to the presence of more submetacentric chromosomes than the other five species. The boxplot illustrates the range and variation of the chromosomal length of six species analyzed (Figure 3). *B. cristata* and *B. prionitis* have larger average chromosomal lengths (ACL) with a wider dispersion (Figure 3). The ACL was similar for *T. erecta* and *T. mysorensis*, as well as for *B. cristata* and *B. prionitis*, indicating a close relationship between the species.

**Figure 3.**
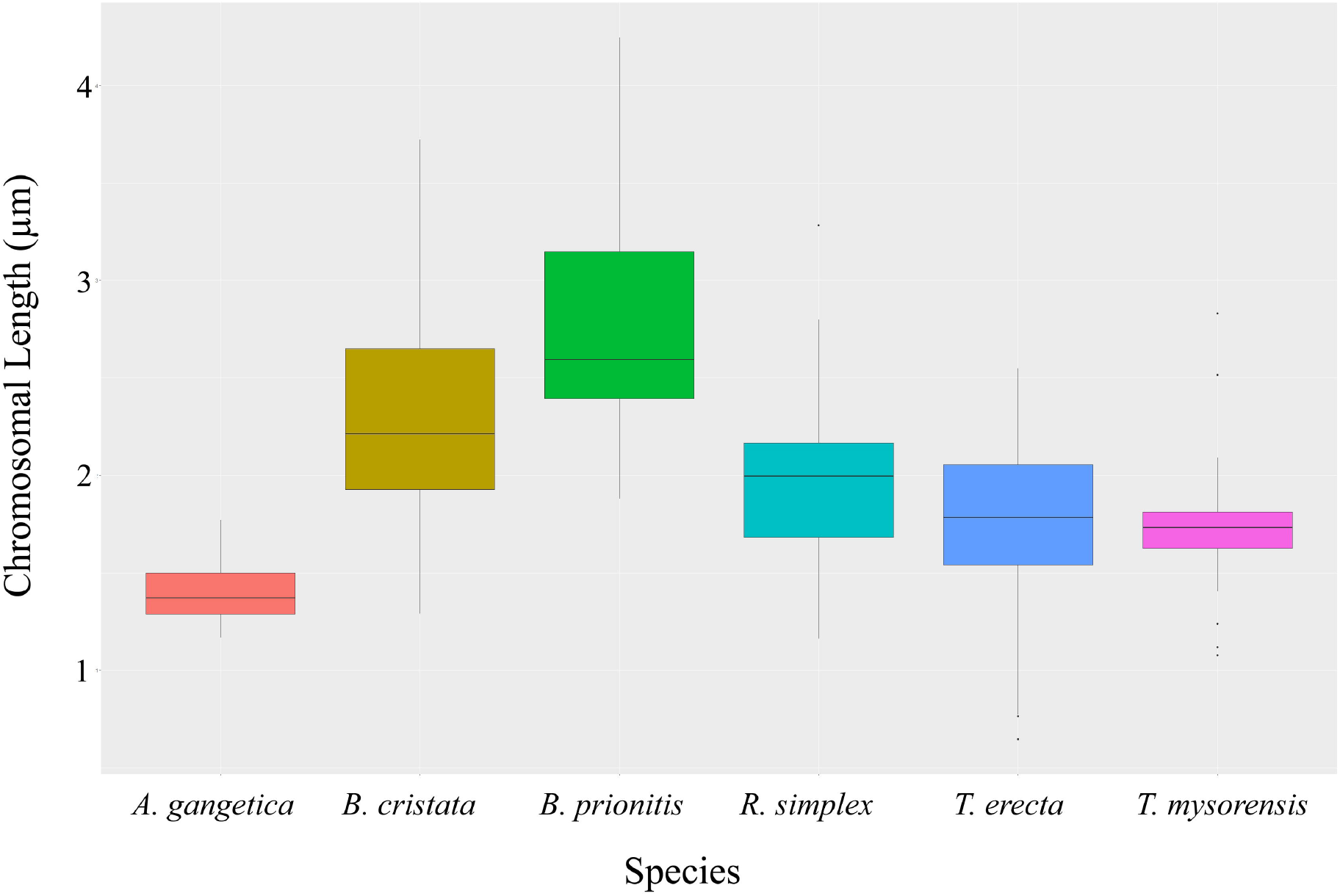
Boxplot representing the chromosomal length diversity among six studied species.

The centromeric formula does not capture the complete degree of asymmetry. That is why several karyotypic symmetric and asymmetric indices have been developed to evaluate karyological variability among closely related species. Among the karyotypic parameters, Syi% and AsK% have preferably displayed a complete negative correlation. The higher Syi% and lower AsK% represent more symmetric karyotype than the lower Syi% and higher AsK%, respectively (Arano 1963; Greilhuber and Speta 1976). The Syi% and AsK% displayed a little variation which was ranging from 82.50–88.17 and 53.14–54.79, respectively (Table 3). Recently, the AI was developed to generate a single number that examined karyotype asymmetry, with higher AI indicating more degrees of karyotypic heterogeneity (Paszko 2006). In this present investigation, the higher AI value of *T. erecta* (2.15) indicates higher karyotype asymmetry among these studied species. According to the foregoing discussion, *T. erecta, R. simplex, B. cristata* and *B. prionitis* were shown to be more advanced than *A. gangetica* and *T. mysorensis* (Table 3). Stebbins (1971) reported that the symmetric karyotype is primitive and the asymmetric is advanced from an evolutionary point of view. From this point, we could say that *T. erecta* showed advanced characteristics among the studied species. Interestingly, the two comparatively primitive species *A. gangetica* and *T. mysorensis* represents racemes types of inflorescence whereas the other studied species shows cyme types of inflorescence (Table 2). We prepared a dendrogram from the karyotypic features. As can be seen in figure 4, all six studied species formed two major clusters. All the species were grouped into the same cluster except *A. gangetica*. The distance was higher between *A. gangetica* and *R. simplex* (1.287) and lower between *B. cristata* and *B. prionitis* (0.021).

**Figure 4.**
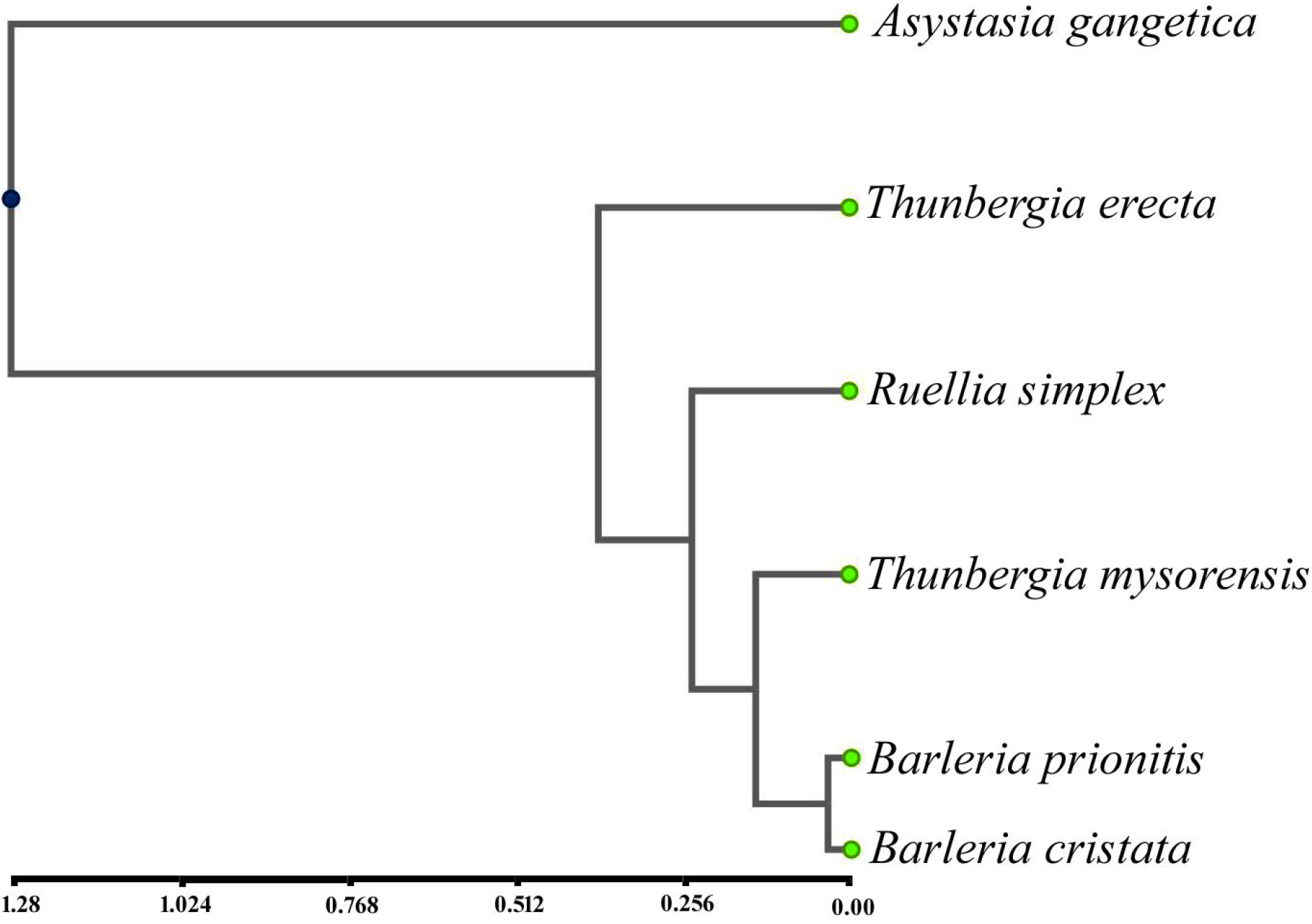
Phylogenetic tree with cluster analysis by UPGMA on the basis of Pearson correlation coefficient of the karyotypes parameters among six examined species.

## Acknowledgements

Authors are grateful to the Department of Botany, University of Dhaka for laboratory facilities.

## Disclosure statement

No potential conflict of interest was reported by the authors.

## Funding information

The authors are grateful to the University of Dhaka for providing financial assistance under the grant from University Grant Commission (UGC), Govt. of the Peoples’ Republic of Bangladesh of fiscal year 2022-2023 (Grant letter no.: 79515-17).

## Notes on contributors

### Nowshin Anjum

MS student, Cytogenetics Laboratory, Department of Botany, University of Dhaka. *Contribution*: carried out the experiments, data analysis and prepared the manuscript.

### Chandan Kumar Dash

Assistant Professor, Cytogenetics Laboratory, Department of Botany, University of Dhaka. *Contribution*: funded, participated in the collection of the plant materials, conceived and designed the study, data analysis and manuscript preparation.

### Syeda Sharmeen Sultana, PhD

Associate Professor, Cytogenetics Laboratory, Department of Botany, University of Dhaka. *Contribution*: funded, conceived and designed the study, supervised experimentation, read and approved the final version of manuscript.

## Notes

### Competing Interest Statement

The authors have declared no competing interest.

