## Supplemental Data 1 for "Karyological diversity among six medicinally important species of Acanthaceae in Bangladesh"

Supplementary Table 1. Length, arm ratio, centromeric index, relative length, centromeric index and centromeric type of mitotic metaphase chromosomes of *Asystasia gangetica*.

| Chrom-osome pair | Long arm (l) µm | Short arm (s) µm | Total length (T) µm | Arm ratio (l/s) | Relative length (RL) % | Centromeric index (CI) % | Centro-meric type (CT) |
|---|---|---|---|---|---|---|---|
| I | 1.11 | 0.66 | 1.77 | 1.68 | 5.25 | 37.25 | sm |
| | 1.11 | 0.62 | 1.74 | 1.78 | 5.15 | 36.00 | sm |
| II | 0.90 | 0.83 | 1.74 | 1.08 | 5.15 | 48.00 | m |
| | 0.80 | 0.76 | 1.56 | 1.05 | 4.63 | 48.89 | m |
| III | 0.78 | 0.77 | 1.55 | 1.01 | 4.59 | 49.78 | m |
| | 0.85 | 0.70 | 1.55 | 1.21 | 4.58 | 45.17 | m |
| IV | 0.88 | 0.60 | 1.48 | 1.47 | 4.39 | 40.54 | m |
| | 0.76 | 0.69 | 1.46 | 1.16 | 4.33 | 47.26 | m |
| V | 0.74 | 0.71 | 1.44 | 1.04 | 4.28 | 49.04 | m |
| | 0.75 | 0.68 | 1.43 | 1.10 | 4.25 | 47.53 | m |
| VI | 0.70 | 0.69 | 1.39 | 1.00 | 4.13 | 49.93 | m |
| | 0.76 | 0.61 | 1.37 | 1.24 | 4.06 | 44.67 | m |
| VII | 0.72 | 0.65 | 1.37 | 1.12 | 4.06 | 47.21 | m |
| | 0.69 | 0.65 | 1.35 | 1.06 | 4.00 | 48.45 | m |
| VIII | 0.74 | 0.61 | 1.35 | 1.20 | 4.00 | 45.36 | m |
| | 0.70 | 0.65 | 1.34 | 1.08 | 3.98 | 48.11 | m |
| IX | 0.72 | 0.61 | 1.33 | 1.18 | 3.96 | 45.83 | m |
| | 0.68 | 0.61 | 1.29 | 1.11 | 3.83 | 47.31 | m |
| X | 0.67 | 0.60 | 1.27 | 1.13 | 3.77 | 46.99 | m |
| | 0.69 | 0.52 | 1.21 | 1.32 | 3.58 | 43.10 | m |
| XI | 0.64 | 0.56 | 1.20 | 1.13 | 3.55 | 46.96 | m |
| | 0.62 | 0.57 | 1.19 | 1.10 | 3.53 | 47.58 | m |
| XII | 0.65 | 0.52 | 1.17 | 1.25 | 3.48 | 44.43 | m |
| | 0.60 | 0.56 | 1.17 | 1.07 | 3.46 | 48.21 | m |
| | | Total = 33.71 µm | | | | | |

m = metacentric chromosome, sm = submetacentric chromosome

Supplementary Table 2. Length, arm ratio, centromeric index, relative length, centromeric index and centromeric type of mitotic metaphase chromosomes of *Barleria cristata*.

| Chrom-osome pair | Long arm (l) µm | Short arm (s) µm | Total length (T) µm | Arm ratio l/s | Relative length (RL) % | Centromeric index (CI) % | Centromeric type (CT) |
|---|---|---|---|---|---|---|---|
| I | 1.96 | 1.76 | 3.72 | 1.11 | 4.01 | 47.39 | m |
| | 1.91 | 1.74 | 3.64 | 1.10 | 3.93 | 47.66 | m |
| II | 1.87 | 1.74 | 3.61 | 1.08 | 3.89 | 48.08 | m |
| | 1.60 | 1.54 | 3.14 | 1.04 | 3.38 | 49.12 | m |
| III | 1.65 | 1.40 | 3.05 | 1.18 | 3.29 | 45.79 | m |
| | 1.65 | 1.33 | 2.98 | 1.25 | 3.21 | 44.52 | m |
| IV | 1.65 | 1.28 | 2.93 | 1.29 | 3.16 | 43.60 | m |
| | 1.58 | 1.26 | 2.84 | 1.26 | 3.06 | 44.25 | m |
| V | 1.36 | 1.33 | 2.69 | 1.03 | 2.90 | 49.35 | m |
| | 1.35 | 1.32 | 2.67 | 1.03 | 2.88 | 49.35 | m |
| VI | 1.33 | 1.31 | 2.64 | 1.02 | 2.84 | 49.47 | m |
| | 1.37 | 1.25 | 2.62 | 1.09 | 2.82 | 47.75 | m |
| VII | 1.27 | 1.26 | 2.53 | 1.01 | 2.73 | 49.86 | m |
| | 1.31 | 1.21 | 2.52 | 1.09 | 2.72 | 47.93 | m |
| VIII | 1.30 | 1.22 | 2.51 | 1.07 | 2.71 | 48.34 | m |
| | 1.24 | 1.17 | 2.41 | 1.07 | 2.60 | 48.41 | m |
| IX | 1.29 | 1.04 | 2.33 | 1.24 | 2.52 | 44.64 | m |
| | 1.28 | 1.01 | 2.29 | 1.26 | 2.47 | 44.24 | m |
| X | 1.20 | 1.07 | 2.27 | 1.12 | 2.45 | 47.09 | m |
| | 1.15 | 1.05 | 2.19 | 1.09 | 2.37 | 47.78 | m |
| XI | 1.42 | 0.81 | 2.23 | 1.77 | 2.40 | 36.14 | sm |
| | 1.39 | 0.81 | 2.19 | 1.72 | 2.37 | 36.71 | sm |
| XII | 1.15 | 1.05 | 2.19 | 1.09 | 2.37 | 47.78 | m |
| | 1.09 | 1.07 | 2.16 | 1.02 | 2.33 | 49.52 | m |
| XIII | 1.19 | 0.97 | 2.16 | 1.22 | 2.33 | 45.02 | m |
| | 1.15 | 0.92 | 2.07 | 1.24 | 2.23 | 44.63 | m |

Table continued….

| Chromosome pair | Long arm (l) µm | Short arm (s) µm | Total length (T) µm | Arm ratio l/s | Relative length (RL) % | Centromeric index (CI) % | Centromeric type (CT) |
|---|---|---|---|---|---|---|---|
| XIV | 1.05 | 0.99 | 2.04 | 1.06 | 2.20 | 48.64 | m |
| | 1.05 | 0.98 | 2.03 | 1.07 | 2.19 | 48.29 | m |
| XV | 1.07 | 0.94 | 2.01 | 1.14 | 2.16 | 46.71 | m |
| | 1.00 | 0.94 | 1.94 | 1.07 | 2.09 | 48.30 | m |
| XVI | 1.01 | 0.87 | 1.89 | 1.16 | 2.04 | 46.32 | m |
| | 0.97 | 0.87 | 1.85 | 1.11 | 1.99 | 47.37 | m |
| XVII | 0.99 | 0.85 | 1.84 | 1.17 | 1.98 | 46.04 | m |
| | 0.88 | 0.84 | 1.72 | 1.04 | 1.86 | 48.99 | m |
| XVIII | 0.84 | 0.76 | 1.60 | 1.10 | 1.73 | 47.62 | m |
| | 0.83 | 0.76 | 1.60 | 1.09 | 1.72 | 47.83 | m |
| XIX | 0.77 | 0.72 | 1.49 | 1.08 | 1.60 | 48.13 | m |
| | 0.76 | 0.72 | 1.47 | 1.06 | 1.59 | 48.58 | m |
| XX | 0.73 | 0.67 | 1.40 | 1.09 | 1.51 | 47.89 | m |
| | 0.71 | 0.58 | 1.29 | 1.22 | 1.39 | 45.01 | m |
| | | Total = 92.76 µm | | | | | |

m = metacentric chromosome, sm = submetacentric chromosome

Supplementary Table 3. Length, arm ratio, centromeric index, relative length, centromeric index and centromeric type of mitotic metaphase chromosomes of *Barleria prionitis*.

| Chrom-osome pair | Long arm (l) µm | Short arm (s) µm | Total length (T) µm | Arm ratio l/s | Relative length (RL) % | Centromeric index (CI) % | Centromeric type (CT) |
|---|---|---|---|---|---|---|---|
| I | 2.28 | 1.97 | 4.25 | 1.16 | 3.76 | 46.31 | m |
| | 2.11 | 2.05 | 4.16 | 1.03 | 3.69 | 49.32 | m |
| II | 1.99 | 1.88 | 3.87 | 1.06 | 3.43 | 48.53 | m |
| | 1.88 | 1.85 | 3.73 | 1.02 | 3.31 | 49.62 | m |
| III | 2.08 | 1.64 | 3.72 | 1.27 | 3.30 | 44.06 | m |
| | 1.98 | 1.71 | 3.69 | 1.16 | 3.27 | 46.33 | m |
| IV | 1.88 | 1.62 | 3.50 | 1.16 | 3.11 | 46.34 | m |
| | 1.91 | 1.51 | 3.42 | 1.26 | 3.03 | 44.17 | m |
| V | 1.82 | 1.37 | 3.19 | 1.33 | 2.83 | 42.86 | m |
| | 1.82 | 1.37 | 3.19 | 1.33 | 2.83 | 42.86 | m |
| VI | 1.60 | 1.54 | 3.13 | 1.04 | 2.78 | 49.09 | m |
| | 1.58 | 1.48 | 3.06 | 1.07 | 2.71 | 48.37 | m |
| VII | 1.91 | 1.23 | 3.13 | 1.56 | 2.78 | 39.09 | sm |
| | 1.82 | 1.17 | 2.99 | 1.56 | 2.65 | 39.05 | sm |
| VIII | 1.60 | 1.42 | 3.02 | 1.12 | 2.68 | 47.17 | m |
| | 1.48 | 1.34 | 2.82 | 1.11 | 2.50 | 47.47 | m |
| IX | 1.42 | 1.34 | 2.76 | 1.06 | 2.45 | 48.45 | m |
| | 1.48 | 1.17 | 2.65 | 1.27 | 2.35 | 44.09 | m |
| X | 1.48 | 1.14 | 2.62 | 1.30 | 2.32 | 43.48 | m |
| | 1.34 | 1.25 | 2.59 | 1.07 | 2.30 | 48.35 | m |
| XI | 1.40 | 1.20 | 2.59 | 1.17 | 2.30 | 46.15 | m |
| | 1.42 | 1.17 | 2.59 | 1.22 | 2.30 | 45.05 | m |
| XII | 1.37 | 1.23 | 2.59 | 1.12 | 2.30 | 47.25 | m |
| | 1.31 | 1.25 | 2.56 | 1.05 | 2.27 | 48.89 | m |
| XIII | 1.40 | 1.14 | 2.54 | 1.23 | 2.25 | 44.94 | m |
| | 1.37 | 1.11 | 2.48 | 1.23 | 2.20 | 44.83 | m |

Table continued….

| Chrom-osome pair | Long arm (l) µm | Short arm (s) µm | Total length (T) µm | Arm ratio l/s | Relative length (RL) % | Centromeric index (CI) % | Centromeric type (CT) |
|---|---|---|---|---|---|---|---|
| XIV | 1.57 | 0.97 | 2.54 | 1.62 | 2.25 | 38.20 | sm |
| | 1.54 | 0.85 | 2.39 | 1.80 | 2.12 | 35.71 | sm |
| XV | 1.34 | 1.08 | 2.42 | 1.24 | 2.15 | 44.71 | m |
| | 1.37 | 1.03 | 2.39 | 1.33 | 2.12 | 42.86 | m |
| XVI | 1.31 | 1.08 | 2.39 | 1.21 | 2.12 | 45.24 | m |
| | 1.34 | 1.03 | 2.36 | 1.31 | 2.10 | 43.37 | m |
| XVII | 1.23 | 1.11 | 2.34 | 1.10 | 2.07 | 47.56 | m |
| | 1.25 | 1.03 | 2.28 | 1.22 | 2.02 | 45.00 | m |
| XVIII | 1.23 | 1.03 | 2.25 | 1.19 | 2.00 | 45.57 | m |
| | 1.28 | 0.94 | 2.22 | 1.36 | 1.97 | 42.31 | m |
| XIX | 1.14 | 1.03 | 2.17 | 1.11 | 1.92 | 47.37 | m |
| | 1.23 | 0.94 | 2.17 | 1.30 | 1.92 | 43.42 | m |
| XX | 1.20 | 0.94 | 2.14 | 1.27 | 1.89 | 44.00 | m |
| | 1.03 | 0.85 | 1.88 | 1.20 | 1.67 | 45.45 | m |
| | | Total = 112.81 µm | | | | | |

m= metacentric chromosome, sm= submetacentric chromosome

Supplementary Table 4. Length, arm ratio, centromeric index, relative length, centromeric index and centromeric type of mitotic metaphase chromosomes of *Ruellia simplex*.

| Chrom-osome pair | Long arm (l) µm | Short arm (s) µm | Total length (T) µm | Arm ratio l/s | Relative length (RL) % | Centromeric index (CI) % | Centromeric type (CT) |
|---|---|---|---|---|---|---|---|
| I | 1.67 | 1.62 | 3.28 | 1.03 | 4.71 | 49.26 | m |
| | 1.49 | 1.31 | 2.80 | 1.14 | 4.01 | 46.65 | m |
| II | 1.72 | 0.88 | 2.60 | 1.95 | 3.73 | 33.87 | sm |
| | 1.44 | 0.94 | 2.37 | 1.53 | 3.41 | 39.47 | sm |
| III | 1.26 | 1.18 | 2.44 | 1.06 | 3.50 | 48.43 | m |
| | 1.15 | 1.08 | 2.22 | 1.06 | 3.19 | 48.44 | m |
| IV | 1.39 | 0.87 | 2.26 | 1.60 | 3.24 | 38.46 | sm |
| | 1.35 | 0.85 | 2.20 | 1.58 | 3.16 | 38.80 | sm |
| V | 1.27 | 0.92 | 2.19 | 1.39 | 3.14 | 41.90 | m |
| | 1.18 | 0.98 | 2.16 | 1.21 | 3.10 | 45.34 | m |
| VI | 1.12 | 1.03 | 2.15 | 1.09 | 3.08 | 47.90 | m |
| | 1.15 | 0.97 | 2.12 | 1.18 | 3.04 | 45.90 | m |
| VII | 1.07 | 1.04 | 2.11 | 1.03 | 3.03 | 49.34 | m |
| | 1.18 | 0.92 | 2.10 | 1.28 | 3.02 | 43.89 | m |
| VIII | 1.08 | 0.97 | 2.05 | 1.11 | 2.94 | 47.46 | m |
| | 1.06 | 0.98 | 2.04 | 1.09 | 2.93 | 47.96 | m |
| IX | 1.06 | 0.98 | 2.03 | 1.08 | 2.92 | 48.12 | m |
| | 1.05 | 0.97 | 2.01 | 1.09 | 2.89 | 47.93 | m |
| X | 1.01 | 0.97 | 1.98 | 1.04 | 2.84 | 49.12 | m |
| | 0.97 | 0.94 | 1.91 | 1.02 | 2.74 | 49.45 | m |
| XI | 0.96 | 0.94 | 1.90 | 1.01 | 2.73 | 49.64 | m |
| | 0.94 | 0.94 | 1.88 | 1.01 | 2.70 | 49.82 | m |
| XII | 0.92 | 0.87 | 1.78 | 1.06 | 2.56 | 48.64 | m |
| | 0.89 | 0.87 | 1.76 | 1.02 | 2.53 | 49.61 | m |
| XIII | 0.92 | 0.84 | 1.76 | 1.09 | 2.52 | 47.83 | m |
| | 0.94 | 0.80 | 1.74 | 1.18 | 2.50 | 45.82 | m |

Table continued….

| Chrom-osome pair | Long arm (l) µm | Short arm (s) µm | Total length (T) µm | Arm ratio l/s | Relative length (RL) % | Centromeric index (CI) % | Centromeric type (CT) |
|---|---|---|---|---|---|---|---|
| XIV | 0.94 | 0.78 | 1.73 | 1.20 | 2.48 | 0.94 | m |
| | 0.80 | 0.75 | 1.55 | 1.06 | 2.22 | 0.80 | m |
| XV | 0.77 | 0.76 | 1.53 | 1.02 | 2.19 | 0.77 | m |
| | 0.76 | 0.76 | 1.51 | 1.00 | 2.17 | 0.76 | m |
| XVI | 0.69 | 0.66 | 1.35 | 1.05 | 1.94 | 0.69 | m |
| | 0.72 | 0.64 | 1.35 | 1.12 | 1.94 | 0.72 | m |
| XVII | 0.68 | 0.62 | 1.31 | 1.09 | 1.87 | 0.68 | m |
| | 0.62 | 0.58 | 1.20 | 1.08 | 1.72 | 0.62 | m |
| XVIII | 0.60 | 0.57 | 1.17 | 1.06 | 1.68 | 0.60 | m |
| | 0.61 | 0.55 | 1.16 | 1.11 | 1.66 | 0.61 | m |
| | | Total = 69.73 µm | | | | | |

m= metacentric chromosome, sm= submetacentric chromosome

Supplementary Table 5. Length, arm ratio, centromeric index, relative length, centromeric index and centromeric type of mitotic metaphase chromosomes of *Thunbergia erecta*.

| Chrom-osome pair | Long arm (l) µm | Short arm (s) µm | Total length (T) µm | Arm ratio l/s | Relative length (RL) % | Centromeric index (CI) % | Centromeric type (CT) |
|---|---|---|---|---|---|---|---|
| I | 1.28 | 1.27 | 2.55 | 1.01 | 2.37 | 49.86 | m |
| | 1.28 | 1.12 | 2.41 | 1.14 | 2.24 | 46.69 | m |
| II | 1.51 | 0.86 | 2.37 | 1.75 | 2.20 | 36.36 | sm |
| | 1.49 | 0.83 | 2.33 | 1.79 | 2.16 | 35.82 | sm |
| III | 1.57 | 0.76 | 2.33 | 2.07 | 2.16 | 32.54 | sm |
| | 1.39 | 0.75 | 2.14 | 1.85 | 1.99 | 35.06 | sm |
| IV | 1.15 | 1.15 | 2.30 | 1.01 | 2.13 | 49.85 | m |
| | 1.08 | 1.05 | 2.12 | 1.03 | 1.97 | 49.35 | m |
| V | 1.28 | 0.92 | 2.21 | 1.39 | 2.05 | 41.82 | m |
| | 1.19 | 0.99 | 2.17 | 1.20 | 2.02 | 45.37 | m |
| VI | 1.06 | 1.04 | 2.10 | 1.02 | 1.95 | 49.50 | m |
| | 1.07 | 1.01 | 2.08 | 1.05 | 1.93 | 48.67 | m |
| VII | 1.09 | 0.99 | 2.08 | 1.11 | 1.93 | 47.49 | m |
| | 1.10 | 0.97 | 2.08 | 1.14 | 1.93 | 46.82 | m |
| VIII | 1.27 | 0.79 | 2.06 | 1.61 | 1.91 | 38.38 | sm |
| | 1.30 | 0.74 | 2.03 | 1.76 | 1.89 | 36.18 | sm |
| IX | 1.06 | 1.01 | 2.07 | 1.06 | 1.92 | 48.66 | m |
| | 1.06 | 0.97 | 2.03 | 1.09 | 1.89 | 47.78 | m |
| X | 1.04 | 0.99 | 2.03 | 1.06 | 1.88 | 48.63 | m |
| | 1.01 | 0.97 | 1.97 | 1.04 | 1.83 | 48.94 | m |
| XI | 1.07 | 0.89 | 1.96 | 1.20 | 1.82 | 45.39 | m |
| | 0.99 | 0.97 | 1.96 | 1.01 | 1.82 | 49.65 | m |
| XII | 0.98 | 0.96 | 1.94 | 1.02 | 1.80 | 49.46 | m |
| | 1.08 | 0.83 | 1.90 | 1.30 | 1.77 | 43.43 | m |
| XIII | 0.99 | 0.90 | 1.88 | 1.10 | 1.75 | 47.60 | m |
| | 1.05 | 0.81 | 1.85 | 1.30 | 1.72 | 43.45 | m |

Table continued….

| Chromosome pair | Long arm (l) µm | Short arm (s) µm | Total length (T) µm | Arm ratio l/s | Relative length (RL) % | Centromeric index (CI) % | Centromeric type (CT) |
|---|---|---|---|---|---|---|---|
| XIV | 0.94 | 0.90 | 1.84 | 1.04 | 1.71 | 49.06 | m |
| | 0.92 | 0.90 | 1.81 | 1.02 | 1.68 | 49.43 | m |
| XV | 1.01 | 0.79 | 1.80 | 1.27 | 1.67 | 44.02 | m |
| | 0.94 | 0.85 | 1.79 | 1.10 | 1.66 | 47.67 | m |
| XVI | 0.90 | 0.89 | 1.78 | 1.01 | 1.66 | 49.81 | m |
| | 0.94 | 0.84 | 1.78 | 1.12 | 1.66 | 47.08 | m |
| XVII | 0.92 | 0.85 | 1.77 | 1.09 | 1.64 | 47.84 | m |
| | 0.90 | 0.87 | 1.76 | 1.03 | 1.64 | 49.21 | m |
| XVIII | 0.93 | 0.80 | 1.73 | 1.17 | 1.61 | 46.18 | m |
| | 0.94 | 0.78 | 1.72 | 1.19 | 1.60 | 45.56 | m |
| XIX | 0.87 | 0.85 | 1.72 | 1.03 | 1.60 | 49.19 | m |
| | 0.94 | 0.77 | 1.71 | 1.22 | 1.59 | 45.12 | m |
| XX | 0.87 | 0.84 | 1.71 | 1.03 | 1.59 | 49.19 | m |
| | 0.85 | 0.84 | 1.69 | 1.02 | 1.57 | 49.59 | m |
| XXI | 0.87 | 0.83 | 1.69 | 1.05 | 1.57 | 48.77 | m |
| | 0.83 | 0.81 | 1.64 | 1.02 | 1.52 | 49.58 | m |
| XXII | 0.83 | 0.80 | 1.63 | 1.04 | 1.51 | 48.94 | m |
| | 0.82 | 0.80 | 1.62 | 1.03 | 1.50 | 49.36 | m |
| XXIII | 0.87 | 0.73 | 1.60 | 1.20 | 1.49 | 45.45 | m |
| | 0.83 | 0.74 | 1.56 | 1.12 | 1.45 | 47.11 | m |
| XXIV | 0.77 | 0.76 | 1.53 | 1.01 | 1.42 | 49.77 | m |
| | 0.76 | 0.75 | 1.51 | 1.02 | 1.41 | 49.54 | m |
| XXV | 0.72 | 0.69 | 1.42 | 1.04 | 1.32 | 49.02 | m |
| | 0.74 | 0.66 | 1.40 | 1.12 | 1.30 | 47.26 | m |
| XXVI | 0.66 | 0.65 | 1.31 | 1.01 | 1.22 | 49.74 | m |
| | 0.68 | 0.62 | 1.31 | 1.09 | 1.21 | 47.87 | m |
| XXVII | 0.67 | 0.63 | 1.30 | 1.05 | 1.21 | 48.66 | m |
| | 0.64 | 0.62 | 1.26 | 1.02 | 1.17 | 49.45 | m |
| XXVIII | 0.63 | 0.62 | 1.26 | 1.01 | 1.17 | 49.72 | m |
| | 0.58 | 0.56 | 1.15 | 1.04 | 1.06 | 49.09 | m |

Table continued….

| Chrom-osome pair | Long arm (l) µm | Short arm (s) µm | Total length (T) µm | Arm ratio l/s | Relative length (RL) % | Centromeric index (CI) % | Centromeric type (CT) |
|---|---|---|---|---|---|---|---|
| XXIX | 0.50 | 0.49 | 0.99 | 1.03 | 0.92 | 49.30 | m |
| | 0.47 | 0.45 | 0.92 | 1.05 | 0.86 | 48.87 | m |
| XXX | 0.44 | 0.41 | 0.85 | 1.08 | 0.79 | 47.97 | m |
| | 0.39 | 0.38 | 0.77 | 1.02 | 0.72 | 49.55 | m |
| XXXI | 0.40 | 0.37 | 0.76 | 1.08 | 0.71 | 48.18 | m |
| | 0.33 | 0.32 | 0.65 | 1.02 | 0.60 | 49.46 | m |
| | | Total = 107.72 µm | | | | | |

m = metacentric chromosome, sm = submetacentric chromosome

Supplementary Table 6. Length, arm ratio, centromeric index, relative length, centromeric index and centromeric type of mitotic metaphase chromosomes of *Thunbergia mysorensis*.

| Chrom-osome pair | Long arm (l) µm | Short arm (s) µm | Total length (T) µm | Arm ratio l/s | Relative length (RL) % | Centromeric index (CI) % | Centromeric type (CT) |
|---|---|---|---|---|---|---|---|
| I | 1.59 | 1.24 | 2.83 | 1.28 | 4.80 | 43.80 | m |
| | 1.43 | 1.08 | 2.51 | 1.32 | 4.27 | 43.09 | m |
| II | 1.05 | 1.04 | 2.09 | 1.01 | 3.55 | 49.83 | m |
| | 1.04 | 0.94 | 1.99 | 1.10 | 3.37 | 47.55 | m |
| III | 1.10 | 0.80 | 1.90 | 1.38 | 3.23 | 41.97 | m |
| | 0.94 | 0.94 | 1.88 | 1.01 | 3.19 | 49.82 | m |
| IV | 1.01 | 0.87 | 1.88 | 1.17 | 3.19 | 46.13 | m |
| | 0.94 | 0.94 | 1.88 | 1.01 | 3.19 | 49.82 | m |
| V | 0.98 | 0.83 | 1.81 | 1.18 | 3.08 | 45.98 | m |
| | 0.91 | 0.90 | 1.81 | 1.01 | 3.08 | 49.81 | m |
| VI | 0.93 | 0.88 | 1.81 | 1.06 | 3.08 | 48.66 | m |
| | 1.08 | 0.73 | 1.81 | 1.49 | 3.08 | 40.23 | m |
| VII | 1.01 | 0.80 | 1.81 | 1.26 | 3.06 | 44.23 | m |
| | 0.90 | 0.87 | 1.78 | 1.03 | 3.02 | 49.22 | m |
| VIII | 1.01 | 0.76 | 1.77 | 1.32 | 3.00 | 43.14 | sm |
| | 0.97 | 0.80 | 1.77 | 1.22 | 3.00 | 45.10 | sm |
| IX | 0.94 | 0.81 | 1.76 | 1.16 | 2.98 | 46.25 | m |
| | 0.87 | 0.83 | 1.71 | 1.05 | 2.90 | 48.78 | m |
| X | 0.87 | 0.83 | 1.70 | 1.06 | 2.89 | 48.57 | m |
| | 0.85 | 0.83 | 1.68 | 1.02 | 2.85 | 49.59 | m |
| XI | 0.94 | 0.72 | 1.66 | 1.30 | 2.82 | 43.51 | m |
| | 0.87 | 0.78 | 1.65 | 1.11 | 2.80 | 47.48 | m |
| XII | 0.86 | 0.78 | 1.65 | 1.10 | 2.79 | 47.68 | m |
| | 0.85 | 0.78 | 1.63 | 1.08 | 2.77 | 48.09 | m |
| XIII | 0.82 | 0.81 | 1.62 | 1.02 | 2.76 | 49.57 | m |
| | 0.83 | 0.79 | 1.62 | 1.05 | 2.76 | 48.72 | m |

Table continued….

| Chrom-osome pair | Long arm (l) µm | Short arm (s) µm | Total length (T) µm | Arm ratio l/s | Relative length (RL) % | Centromeric index (CI) % | Centromeric type (CT) |
|---|---|---|---|---|---|---|---|
| XIV | 0.81 | 0.81 | 1.62 | 1.01 | 2.75 | 49.79 | m |
| | 0.84 | 0.69 | 1.53 | 1.21 | 2.60 | 45.25 | m |
| XV | 0.77 | 0.69 | 1.47 | 1.11 | 2.49 | 47.39 | m |
| | 0.76 | 0.69 | 1.45 | 1.09 | 2.46 | 47.85 | m |
| XVI | 0.72 | 0.69 | 1.40 | 1.04 | 2.38 | 49.01 | m |
| | 0.63 | 0.60 | 1.24 | 1.05 | 2.10 | 48.88 | m |
| XVII | 0.60 | 0.52 | 1.12 | 1.15 | 1.90 | 46.58 | m |
| | 0.56 | 0.51 | 1.08 | 1.09 | 1.83 | 47.74 | m |
| | | Total = 58.93 µm | | | | | |

m= metacentric chromosome, sm= submetacentric chromosome
